# The Effect of Dissociation between Proprioception and Vision on Perception and Grip Force Control in a Stiffness Judgment Task

**DOI:** 10.1101/252536

**Authors:** Stephanie Hu, Raz Leib, Ilana Nisky

## Abstract

Our sensorimotor system estimates stiffness to form stiffness perception, such as for choosing a ripe fruit, and to generate actions, such as to adjust grip force to avoid slippage of a scalpel during surgery. We examined how temporal manipulation of the haptic and visual feedback affect stiffness perception and grip force adjustment during a stiffness discrimination task. We used delayed force feedback and delayed visual feedback to break the natural relations between these modalities when participants tried to choose the harder spring between pairs of springs. We found that visual delay caused participants to slightly overestimate stiffness while force feedback delay caused a mixed effect on perception; for some it caused underestimation and for some overestimation of stiffness. Interestingly and in contrast to previous findings without vision, we found that participants increased the magnitude of their applied grip force for all conditions. We propose a model that suggests that this increase was a result of coupling the grip force adjustment to their proprioceptive hand position, which was the only modality which we could not delay. Our findings shed light on how the sensorimotor system combines information from different sensory modalities for perception and action. These results are important for the design of improved teleoperation systems that suffer from unavoidable delays.

## I. Introduction

When interacting with a mechanical spring, the central nervous system combined haptic, proprioceptive, and visual information to form a perception of stiffness [1-3]. In particular, it has been proposed that visual and haptic information are optimally integrated such that the stiffness estimation is a weighted average of the two modalities. Weights are assigned according to the variance of each input: the sense with less noise contributes more to the combined percept [1]. Visual information often dominates in the perception of object properties like size and shape, a phenomenon known as “visual capture” [4, 5]. Interestingly, studies have shown that vision alone can be reliably used to estimate the softness of an object, and that when haptic and visual information are combined, the resulting integration is biased towards vision [3]. Yet, the contribution of visual information to the way people interact with the object during softness discrimination is yet unknown.

Psychophysical studies showed that manipulating both haptic and visual input can influence perceived stiffness [2, 6]. DiLuca et al. (2011) demonstrated that delaying visual feedback causes overestimation of stiffness due to the mismatch between the amount of displacement sensed proprioceptively versus the amount of displacement acknowledged visually. Their experiment also supported previous research showing that participants underestimate stiffness of springs with delayed force feedback [7-9].

Stiffness is also estimated in order to predict load force during interactions with springs. During dynamic object manipulation, where we grasp an object that is attached to a spring, we experience load force generated by the lengthening or shortening of the spring. To avoid object slipping from our grasp, we form an internal model that anticipates the load force during movement and generates appropriate grip force that slightly lead the load force [10]. During initial interaction with the object, the spring's stiffness is unknown, and we use excessive grip force to avoid object slipping; however, after a few interactions the motor system can predict the load force and reduce the grip force magnitude to avoid fatigue [11].

Leib et al. (2015) examined stiffness perception and grip force adjustment on the probing tool during interaction with a virtual elastic force field. While there was a discrepancy in the subjects’ perception responses between the delayed and non-delayed force fields, their grip force characteristics were similar when interacting with each field. Grip force magnitude decreased between early and late probes and the grip force trajectory was shifted in time to temporally align with the load force trajectory. This suggests that separate neural streams are responsible for perception and action, and that while the action stream is able to form an accurate representation of stiffness and time delays, the perception stream cannot [9].

The objective of this study is to examine the effects of visual feedback on perception and grip force adjustment during a stiffness discrimination task. We aimed to test if the discrepancy between stiffness perception and grip-force modulation exists when force feedback is delayed but spring displacement remains visually aligned with proprioceptive signals. In addition we tested how only visual delay and a combination of visual and force feedback delay affect action and perception. Our results indicate that the presence of visual information breaks the coupling between load force and grip force in force-delayed springs, suggesting that there is a bias towards visual cues not only in perception, but also action.

## II. Materials & Methods

### A. Participants & Setup

Ten participants (five females and five males, ages 18-26, all right handed) participated in the experiment after signing a consent form. The study was approved by the Human Subjects Research Committee of Ben-Gurion University of the Negev, Beer-Sheva, Israel.

Participants sat in front of a virtual reality setup while holding the end-point of a robotic device as depicted in (Fig. 1A). We instructed participants to hold the device using their thumb and index fingers in a way that resemble holding a tool. We equipped the end point of the haptic device (Phantom Desktop; Sensable Technologies) with a grasp fixture with a force sensor (Nano17; ATI Technologies) to record the grip forces that were applied during the experiment. The position of the tool (as measured based on the readouts of the haptic device encoders) was recorded at 1 KHz and used to calculate, in real time, a force feedback that was applied via the haptic device. Grip force was recorded via the force sensor at 1 KHz. Participants looked at a semisilvered mirror showing the projection of an LCD screen (60 Hz refresh rate) placed horizontally above it. An opaque screen was fixed under the mirror to block the vision of the hand. The position of each participant’s hand was displayed as a blue sphere. In addition, a cloth was draped over the participant’s shoulders as a second measure to block actual visual feedback.

**Figure 1.**
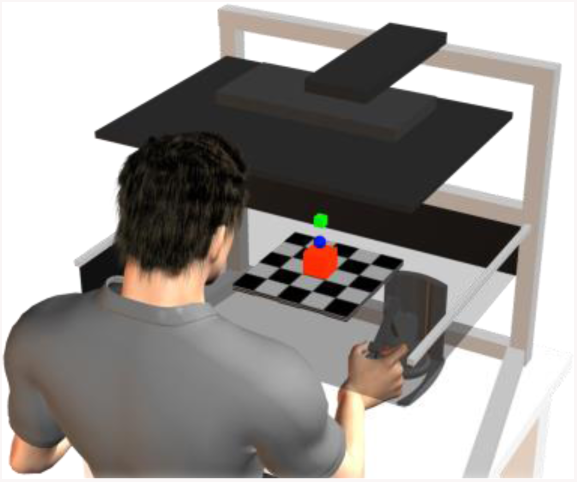
Experimental setup. Participant moved a blue sphere used to compress and release a spring displayed as rectangular prism. Participants were asked to examine a blue and red springs and decide which one is stiffer by indicating the stiffer spring color. To switch between springs the participant elevated his hand to reach a green cube. Participant held the stylus of the haptic device using their thumb and index fingers.

Participants interacted with two different springs during each trial and reported which one they perceived to be stiffer. The springs were displayed as rectangular prisms sitting on a checkerboard-patterned floor, and were one of two colors: blue or red. Participants could switch between the two springs by placing the sphere over a virtual green button, represented on screen as a cube fixed at a distance of 6 cm above the resting position of the spring. We did not restrict the number of times participants were allowed to switch between springs or the number of probing movements they were allowed to perform during interaction with each individual spring; however, we did ask for participants to continuously probe the spring without breaking contact with it to the best of their abilities. Participants moved the haptic device up and down when probing the artificial spring to simulate pressing down on a real spring. When ready to answer, they verbally responded the color of the spring that they judged to be stiffer, and the experimenter would enter the keyboard key that was marked with the corresponding color. Pressing of the keyboard caused the spring to disappear from the screen, indicating the start of a new trial. Participants initiated the trial by moving their cursor over the green button.

### B. Experiment Protocol

During the probing interaction, the applied forces, F(t), were calculated according to 
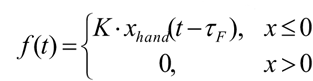
 where K [N/m] is the stiffness, and τ_F_ [ms] is the force feedback delay. Participants were also given visual feedback about their hand position according to 
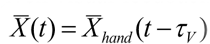
 where 
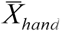
 is the end point of the haptic device, τ_V_ [ms] is the is dback delay. We used different values for τ_V_ and τ_F_ that set four conditions: no delay, τ_V_=0, τ_F_=0; force delay only, τ_V_=0, τ_F_=50, visual delay only, τ_V_=50, τ_F_=0, and both visual and force delay, τ_V_=50, τ_F_=50. The magnitude of force delay is the same as in previous experiments conducted by two of the authors [7, 9]; the visual delay was then chosen to be 50ms so that force and visual feedback would be aligned during the both delayed condition.

Each trial consisted of a pair of virtual elastic springs, a *standard* and a *comparison*, presented to the subject as described in the previous section; both the order in which they were presented and the color assigned to each spring was random. The simulated stiffness of the standard spring was held constant at *K_S_* = 85 N/m in all the trials, though the stimulus varied between trials with regards to visual and/or force delay. In the force delay and both delay conditions, the force lagged the displacement of the spring by 50ms; in the no delay and visual delay conditions, no haptic delay was introduced (except from the 2-3ms it takes for our system to calculate the force feedback that was inherent to all the virtual springs). Similarly, in the visual delay and both delay conditions, we introduced a 50ms latency in the rendering on screen. The inherent visual delay was 16ms for all springs, so the 50ms in the visual delay and both delay conditions was added on top of that. The comparison springs took on simulated stiffness levels between 40-130 N/m, spaced at intervals of 10 N/m; these springs were all non-delayed. To prevent subjects from reaching the force saturation levels of the haptic device by pressing to far down on the spring, a beep was played when the deformation reached 5 cm, without regards to the spring type or stimulus.

Each subject participated in 340 trials split evenly across two days (170 trials per day). The process took about 40 minutes to complete the 170 trials each day; the participants were allowed to ask to take a break at any point in the experiment, as many times as they wanted and for as long as they wanted. The first ten trials on each day were practice trials in which participants familiarized themselves with the system. After the training session, the subjects completed an additional 160 trials each day. The 160 trials each day were split evenly between the four stimulus conditions: no delay, force delay, visual delay, and both force and visual delay, and the different conditions were mixed and not blocked. In total, there were 40 different standard-comparison spring combinations, and each combination was presented to the subjects 4 times per day, for a total of 8 across both days. The order in which the spring pairs was presented was pseudorandomly predetermined across all trials; within each trial, the order in which the standard and comparison springs were presented was random. Participants did not receive feedback about their answers, even during the training trials.

### C. Data Analysis

*Grip force modulation analysis*. Most of the data analysis was completed according to procedures discussed in length in [9]. We analyzed the grip force adjustment by examining the position, load, and grip force trajectories (Fig. 2A) and the trajectories in the grip force-load force plane (Fig. 2B) [9, 12]. We analyzed grip force modulation by examining the data during the first interaction with the spring, when participants were unaware of the standard spring type, and during the last probing movement before they decided about which spring is stiffer. For each trial, we fitted a regression line to the data in the grip force-load force plane for the first and last probing movements (see example in Fig. 2B). We extracted the slope and intercept values from each regression line. According to [12], when the slope of the regression lines are similar between trajectories, the change in intercept values indicates a change in grip force magnitude. For example, an increase in the intercept value between a regression fitted to the data of the first and the last probing movement indicates an increase in grip force magnitude between the two movements. In addition, we calculated the temporal lag between the grip force and load force signals by computing the lag between the peaks of the signals, and verified the results by calculating the cross-correlation function between the two signals [9].

**Figure 2.**
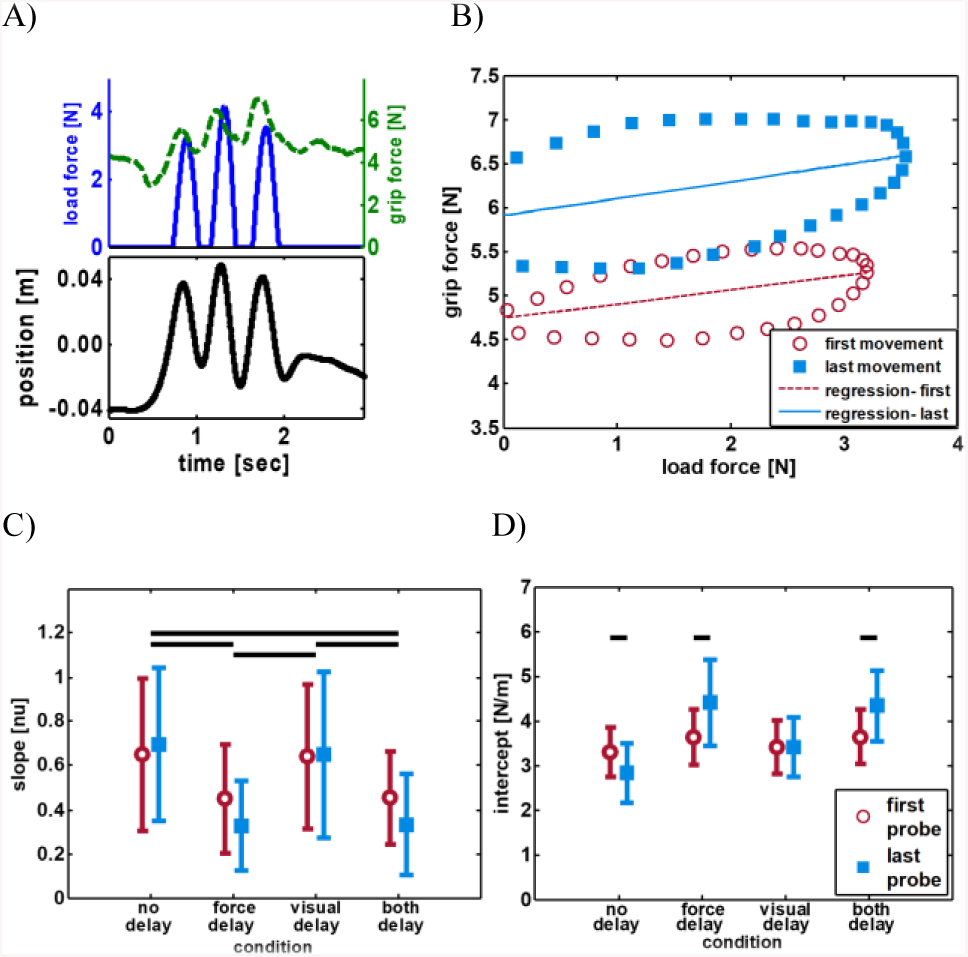
A) Example for probing movement of a standard spring where both visual and force feedback were delayed. Upper panel shows an example for the grip force (dashed green line) and load force (solid blue line) trajectories. Bottom panel shows the tool position. Light blue shaded squares mark the first and last probing movements data we used for the regression analysis. B) First (red empty square) and last (blue filled square) probing movement trajectories from A plotted in the grip force-load force plane. Regression lines fitted to the data of each movement are indicated by the color of the trajectory. C) General analysis for slope value extracted from the regression lines for the four conditions we in the experiment separated to the values extracted from the first probing movement and from the last probing movement. Horizontal lines indicate a statistically significant difference (p<0.05). D) Same as in C but for the intercept value.

*Perceptual responses analysis*. We used the answers of the participants to calculate the probability of a subject to answer “comparison spring had higher level of stiffness” as a function of the difference between the levels of stiffness of the comparison and the standard springs. We used the psignifit 2.5.6 toolbox [13] to fit logistic psychometric curves to the probabilities, and extracted the point of subjective equality (PSE) and the just noticeable difference (JND) as described in detail in [7]. The PSE indicates the difference in stiffness at which there is a 0.5 probability of the subject answering that the comparison spring is stiffer; for example, a positive PSE value of 25N/m means that the participant judged the stiffness of the standard spring to be equal to a comparison spring with a stiffness of 110N/m. The JND describes how much the stiffness levels of the springs must differ for the subject to notice a discrepancy.

*Statistical analysis*. For the standard spring, to test the effect of the feedback delay and probing movement on the slope and intercept of the regression lines fitted to the grip-force load force trajectories, we fitted two separate two-way repeated-measures ANOVA. The *slope* or *intercept* values were the dependent variable, and *condition* (no delay, force delay, visual delay, both delay) and *probing movement* (first and last) were the within-subject independent factors. A similar model was fitted to test the effect of the four conditions and the probing movement on the grip force-load force lag. To test the effect of the four conditions on stiffness perception we fitted two one-way repeated-measures ANOVA models with *PSE* and *JND* as the dependent variable and *condition* as an independent factor. Post hoc tests with Bonferroni correction for multiple comparisons were performed when factors had more than two levels. Statistical significance was determined at the 0.05 threshold in all tests.

## III. Results

In this study, we found that introducing delay to the force feedback, the visual feedback and a combination of them increases the grip force magnitude. During interaction with non-delayed virtual springs, participants reduced their grip force between the initial and the final interaction with the spring. When we delayed the visual feedback, we did not observe such decrease. Introducing force delay significantly increased the magnitude of the grip force. In contrast to this consistent increase in grip force magnitude, we observed a mix effect on perception; delayed visual feedback caused an overestimation and delayed force feedback can cause either over-or-under estimation of stiffness.

### Delayed force feedback increase the grip force magnitude

Grip force magnitude increased in conditions involving force delay. During multiple interactions with the standard spring, participants changed their applied grip force. When force feedback was synchronized with the hand position, i.e. no delay and visual delay conditions, we observed a decrease or no change in magnitude of the grip force, respectively. However, when force feedback was delayed, i.e. during the force delay and both delay conditions, we observed that participants increased their grip force. An example for an interaction with a standard spring with force feedback delay is depicted in Fig. 2A. In this example, the grip force increased between the first and last probing while the load force remained similar. These observations were supported by the analysis of the regression coefficient that were fitted to the grip force-load force trajectories. Using this analysis we observed that the intercept of the regression lines increased between the first and last probing movement while the slope remained similar between these movements (Fig. 2B).

These observations are representative of the entire population. For the fitted regression slope value (Fig. 2C), we found that the slope did not change between the first and last probing movement (F_1,9_=2.62, p=0.14), but was different across the four delay conditions (F_3,27_=23.9, p<0.001). Post hoc analysis showed that this effect was due to differences between the no delay and force delay conditions (t_9_=5.1,p=0.003), the no delay and both delay (t_9_=5.1,p=0.003), the visual delay and force delay (t_9_=4.8,p=0.006) and visual delay and both delay (t_9_=4.6,p=0.007). In addition, we did not find statistically significant interaction between the probing movement factor and condition factor (F_3,27_=2.78, p=0.06). This means that the slope of the regression line did not differ between the first and last probing movements within a single condition.

For the intercept value (Fig. 2D), we found statistically significant effect of the probing factor (F1,9=5.92, p=0.038), the conditions factor (F_3,27_=33.0, p<0.001), and their interaction (F_3,27_=42.5, p<0.001). Post-hoc analysis showed that the intercept value was smaller between the first and last probing movement for the no delay condition (t_9_=6.75, p<0.001), and was higher for the force delay condition (t_9_=4.32, p=0.002) and for the both delay condition (t_9_=4.32, p=0.002). Taken together, the lack of change in slope value and the change in intercept value mean that the magnitude of the grip force reduced between the first and last probing movements for the no delay condition, did not change for the visual delay condition and increased for the force delay and both delay conditions. These results suggest that both delayed force feedback and delayed visual feedback increase the grip force magnitude, however, the delayed force feedback has a larger effect on the grip force compared with the delayed visual feedback.

### In conditions involving force delay, grip force signal does not adjust to load force signal

On average, the grip force led the load force by 54.1 ± 23.7 ms in the first probe and 41.8 ± 18.0 ms when interacting with the standard no-delay spring, reflecting the predictive nature of grip force adjustments to load force. Introducing delay causes the load force to lag hand position, so in order to prevent slippage of the haptic device, grip force magnitude either needs to be increased or its trajectory needs to be shifted to align with the load force.

Previous studies without visual feedback showed that grip force is aligned with load force when probing a linear elastic force field, and it is shifted in time in the presence of force feedback delay [9]. In contrast with these results, our results show that when visual feedback is introduced, grip force no longer shifts towards the load force in the presence of force feedback delay (F_1,9_=2.77, p=0.13) (Fig. 3A). In the force-delayed spring, the grip force led the load force by 78.2 ± 20.6 ms in the first probe and 74.5 ± 17.5 ms in the last probe; similarly, in the both-delayed spring, the grip force led the load force by 75.6 ± 18.1 ms in the first probe and 69.3 ± 14.6 ms in the last probe. Moreover, there was no interaction between the probing movement and condition terms (F_3,27_=1.45, p=0.24): none of the conditions showed a shift of the grip force between the first and last probes.

**Figure 3.**
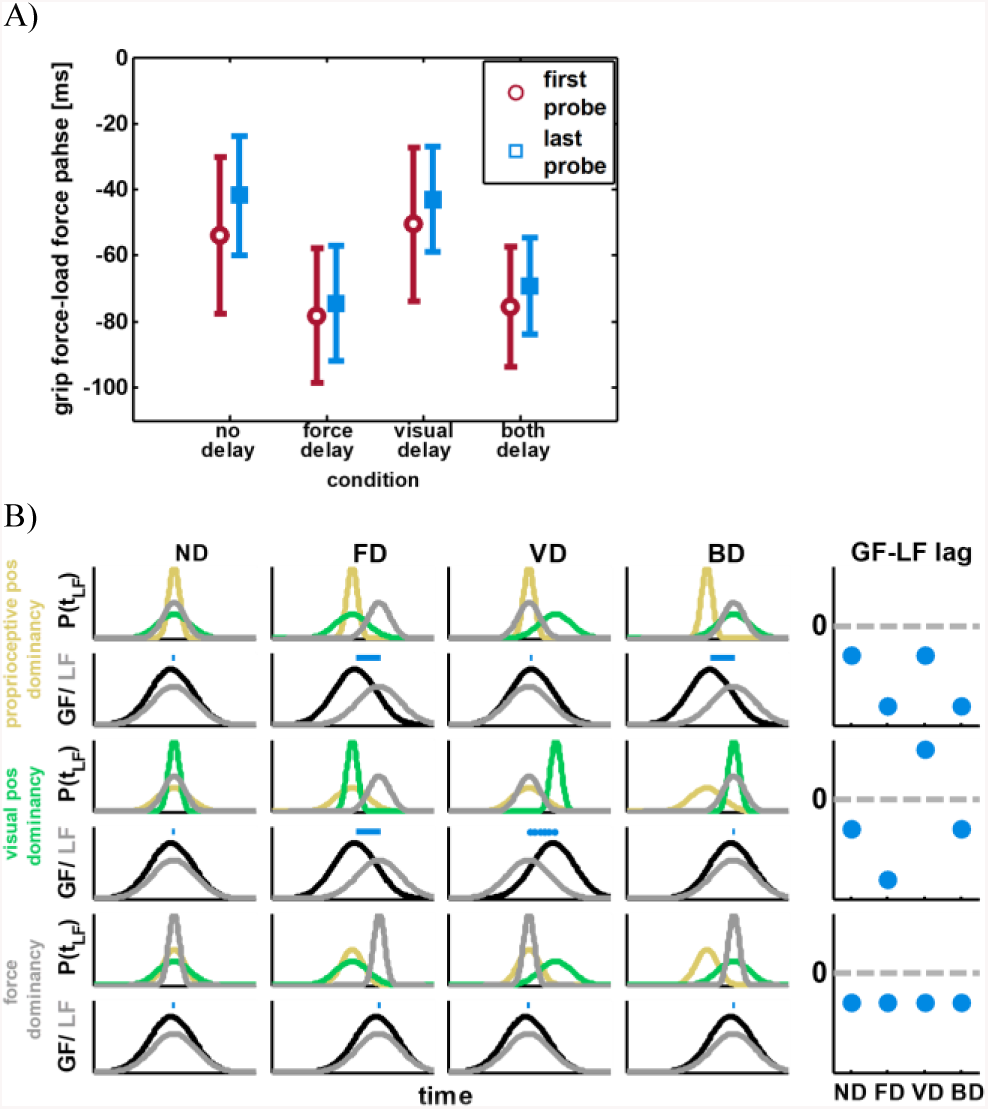
A) Lag analysis between grip force and load force. For each condition we plotted the temporal lag between the peak of the grip force and the peak of the load force during the first probing movement (red empty circle) and during the last probing movement (blue filled squares). For conditions in which force feedback was delayed we observed an increase in lag value compared to conditions with force feedback that was synchronized with tool position. This increased lag was similar between the first and last probing movements. B)Simulation results. We simulated the probability for the timing of peak load force during the last probing movement as a weighted sum of the probability from three sensory modalities: proprioceptive position (yellow line), visual position (green line), and sensed force (gray line). The first two rows of panels depict proprioceptive position dominancy, the middle two rows depict visual position dominancy, and bottom two rows depict sensed force dominancy. In the second row of panels we illustrate for each scenario and condition the grip force (black line) and load force (gray line) trajectories. Blue horizontal lines indicate the lag between grip force and load force. Solid blue line indicate negative lag and dotted blue line indicate positive lag. We summarize the value of the resulted lag in the right column for each scenario. Comparing the simulation results to the actual results of the lag during last probing movement (A, blue squares) we observed that grip force was coupled with the proprioceptive position modality (first two rows).

The grip force-load force lag did not differ between the first and last probes within a given condition, but post hoc analyses revealed that the lag varied between different conditions.

Specifically, we found that there were differences between the conditions involving no delay and force delay (t_19_=8.7, p<0.001), no delay and both delay (t_9_=8.70, p<0.001), visual delay and force delay (t_9_=9.19, p<0.001), and visual delay and both delay (t_9_=8.27, p<0.001). Thus, the lag between the load force and grip force increased by approximately the same amount in both the first and last probing movements whenever the force feedback was delayed.

To explain the results of the grip force timing, we assumed that the motor system can estimate the timing of the peak load force from three sources – the proprioceptive position of the hand, the visual position of the hand, and the sensed force. Under this assumption, we simulated the probability for the timing of the peak load force during the last probing movement as a weighted sum of the probability from these sensory modalities (Fig. 3B). In each simulation, we considered that the weight of one of the modalities as larger than the other two by setting this modality to have higher probability to accurately estimate the peak load force. Since the actual estimation is a weighted sum of all three probability functions, it will be closer in value to the estimation with the higher probability. After simulating the probabilities functions for each of the four experimental conditions we used in the experiment, we calculated an estimation for peak load force time, and simulated a trajectory of the grip force such that its peak slightly led the estimated load force peak. We found that the results of the lag between grip force and actual load force best fit a simulation in which grip force is coupled with the proprioceptive hand position.

### Perception is biased in delayed conditions

All the participants, except for 1, slightly overestimated the stiffness of the standard spring in the presence of visual delay. Delaying the force feedback caused a mixed effect; some participants underestimated the stiffness (6 subjects) and some overestimated the stiffness (4 subjects). An example for a participant who underestimated the standard spring stiffness with force delay is depicted in Fig. 4A. We observed that the PSE value for the both delay condition was between the values of the force delay and the visual delay values for 6 out of 10 participants (Fig. 4B). However, since participants did not exhibit any consistent effect between conditions, we did not find a statistically significant effect of the delay conditions on stiffness perception (F_3,27_=1.08, p=0.37). Similarly, the delay conditions did not have any statistically significant effects on the JND (Fig. 4C) (F_3,27_=1.78, p=0.174).

**Figure 4.**
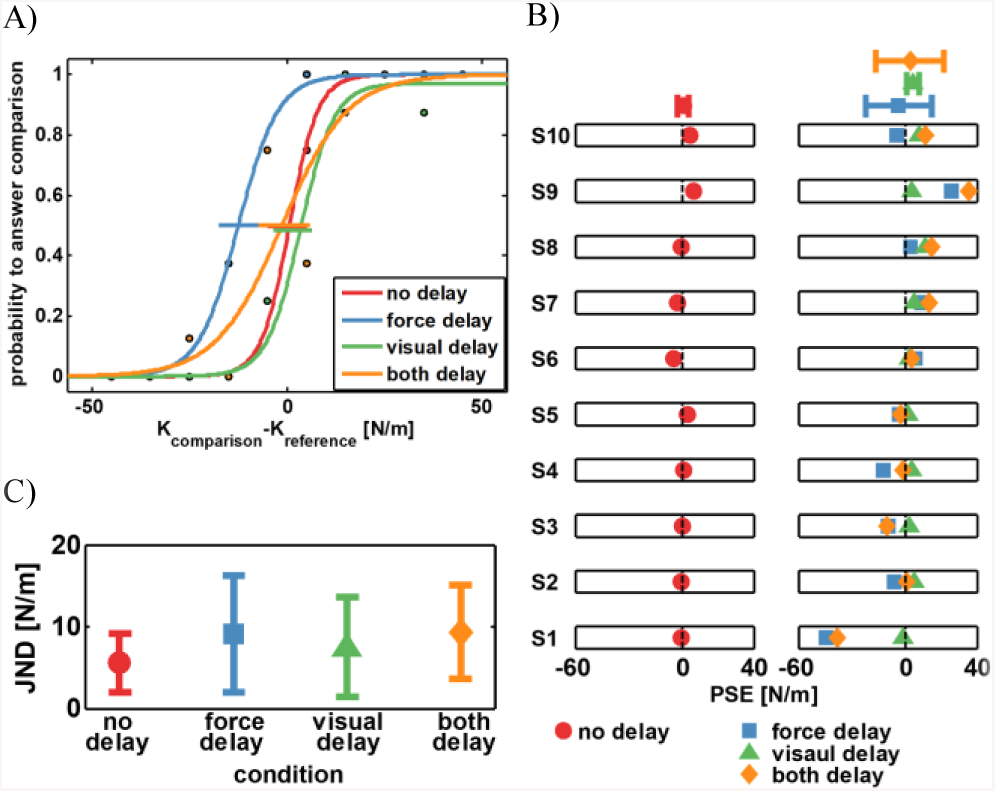
A) An example for psychometric curves fitted to participant’s responses for the four conditions: no delay (red), force delay (blue), visual delay (green), and both delay (orange). In this example, the participant overestimated the standard spring with visual delay and underestimate the spring with delayed force feedback. When both modalities were delayed the participant showed a weighted response. B) individual PSE value. Each raw represent the PSE values of one participant for all four conditions. The four conditions are represnted by distinct color and shape: red circle (no delay), blue square (force delay), green triangle (visual delay), orange dimond (both delay). Points above the individual results indicate mean PSE value for each condition and error bars represent STD value. Vertical black dashed lines mark the zero PSE value. C) Each point represent mean JND value. Error bars indicate STD value.

## IV. Discussion

In our experiment, we examined how delayed force and visual feedback affect stiffness perception and grip force modulation during interaction with a virtual linear elastic spring. We tested four conditions: no delay, force delay, visual delay, and both force and visual delay. Our results demonstrate that visual feedback that is misaligned in time with either force feedback, proprioception, or both during stiffness discrimination increases grip force magnitude and decouples grip force-load force alignment. Thus, adding temporally misaligned visual information increased the uncertainty of the participants about the time when the load force will increase during interaction with the spring, resulting in increasing the grip force to avoid slippage.

In line with our previous results, in the no delay springs, the participants adjusted and reduced the magnitude of the grip force between early and late probing movements in accordance with the stiffness of the spring [9]. However, in the case of delayed force feedback, in contrast to [9], we observed an increase in grip force magnitude during late probing movements. We reason this difference in results between the two studies is due to the changes in the probability function of the timing of the estimated load force that is used for the anticipatory adjustment of grip force with load force. According to the maximum-likelihood integration hypothesis [1], the estimation is biased towards the more certain information channel. In our case, proprioceptive position, visual position, and sensed force contributed to the estimation of the trajectory of the load force: Our results indicate that participants relied on the proprioception, and coupled the predictive grip force with the hand position. Coupling the grip force with actual tool position when the load force is delayed can result in the haptic device slipping, and to overcome this, participants increased the magnitude of the grip force to increase grip force safety margin [11]. Interestingly, the contrast to the study by [9] where the grip force was coupled with the load force suggests that introduced the visual feedback changed the control of grip force.

Based on our results, we argue that the addition of visual feedback to the task of probing a linear elastic spring prevented forming an accurate representation of both stiffness and time delay. Leib et. al. (2015) showed that perception of stiffness was able to help adjust the grip force in accordance with the load force. With the use of this internal representation, the grip force adjustment process may be a result of interaction between feed-forward and feedback control mechanisms. In such a view, the feed-forward mechanism is responsible for generating the grip force in accordance with a predicted load force, and the feedback mechanism is responsible for adjusting the internal representation of dynamic properties, such as the stiffness, based on the sensory input [12, 14]. The latter requires combination between position and force to estimate stiffness[15].The misalignment of these signals, then, results in biased perception and inappropriate grip force adjustment.

It is noteworthy to state that our simulations, which lead us to this explanation, relate to the grip force control mechanism and not to the mechanism of perceptual estimation of stiffness. Future studies may focus on the mechanism for forming stiffness perception based on information that are received from different modalities and its link or lack of to the increased weight of the proprioceptive position for grip force control we showed here.

There are several open questions in our results. First, the participants could not anticipate the condition in each trial, and therefore, in conditions with force delay we expected an increase of roughly 50 ms in grip force-load force lag in the first interaction. However, we observed a smaller increase in the lag. One possible explanation is that the dominant peaks in the grip force were of a reactive nature and not of a predictive nature [16]. In such case, the calculation of the lag would show smaller value than 50 ms.

Second, it is noteworthy that in our experimental paradigm we did not examine how the virtual force fields are perceived compared with actual springs. A possible explanation for the lack of grip force adaptation is the fact that virtual force fields are inherently perceived to be stiffer than real springs, leading to a less optimized modulation of grip force than “normal” springs. This however does not explain the discrepancy between our current study and [9], where using a very similar paradigm adaptation of grip force to the correct timing and the correct stiffness level was reported. The only difference between our study and [9] is the added visual feedback.

Third, our results for the perceptual responses show that introducing visual delay biases stiffness perception. However, in contrast to previous studies [2], visual delay caused only slight overestimation of the spring stiffness, if any. One noticeable difference between the two experiments is the fact that we used a small value for the visual delay. Although this value is presumably above the noticeable value [17] and may trigger an implicit effect on participants' perception, it is likely that increasing the visual delay will increase the impact of the delay on perception, resulting in larger overestimation of stiffness. We plan to repeat this study using a larger value for the visual delay, keeping the force delay at 50 ms.

## V. Conclusion

We studied how visual and force delay affect stiffness perception and grip force modulation when performing tool-mediated interaction with a virtual spring. The addition of visual feedback decouples the alignment of grip force and load force such that grip force is modulated according the actual position of the tool, rather than the load force signals. Understanding how the nervous system integrates, perceives, and responds to information from its environment is critical for the development of efficient human-machine interfaces that have inherent and unavoidable delays in force and visual feedback. This is applicable to a wide range of fields, including rehabilitation, teleoperation, and surgery [18].

